# Experimental flooding impacts soil biogeochemistry but not aboveground vegetation in a coastal forest

**DOI:** 10.1101/2025.05.09.653188

**Authors:** Peter Regier, Ben Bond-Lamberty, Nicholas Ward, Vanessa Bailey, Roberta Peixoto Bittencourt, Fausto Machado-Silva, Nate Mcdowell, Kendalynn Morris, Allison Myers-Pigg, Steph Pennington, Mizanur Rahman, Roy Rich, Richard W. Smith, Stephanie Wilson, Stella C. Woodard, Alice Stearns, Donnie Day, Kennedy Doro, Efemena Emmanuel, Olawale Ogunsola, Kaizad Patel, Pat Megonigal

**Affiliations:** Marine and Coastal Research Laboratory, Pacific Northwest National Laboratory, Sequim, WA; Pacific Northwest National Laboratory, Richland, WA; University of Toledo, Toledo, OH; Smithsonian Environmental Research Center, Edgewater, MD; Global Aquatic Research LLC, Sodus, NY

**Author notes:** Now at Department of Forestry and Environmental Science, Shahjalal University of Science and Technology, Sylhet, Bangladesh (Mizanur Rahman). **Corresponding Author:** Peter Regier. **Author Contributions:** Conceptualization - VB, PM, AM-P, SP, PR, RS, SCW Data Curation - FMS, BP, SP, MR, PR, RR, SJW Formal Analysis - KM, PR Funding acquisition - VB, BB-L, PM, NW Investigation - BB-L, DD, KD, EE, FMS, NM, PM, AM-P, OO, KP, RP, PR, RR, NW Methodology - FMS, NM, PM, KM, AM-P, RP, SP, MR, PR, RR, RS, AS, NW, SJW, SCW Project administration - VB, PM Resources - PM, PR, RS, AS, SJW, SCW Software - PR Supervision - VB, BB-L, NM, PM, NW Validation - BB-L, KM, MR, PR Visualization - PR Writing – original draft - PR, BB-L, NW Writing – review & editing - All authors. **Competing Interest Statement:** The authors declare they have no competing interests.

**Keywords:** Coastal forests, flooding, ecosystem change, soil biogeochemistry, tree physiology

## Abstract

Rising sea levels and intensifying storms increase flooding pressure on coastal forests, triggering tree mortality, ecosystem transitions, and changes to the coastal carbon cycle. However, the mechanisms that drive coastal forest mortality remain elusive due to the complex interplay between belowground and aboveground processes during flooding disturbances and limitations of observations typically reported in coastal forest mortality studies. We used an ecosystem-scale manipulation to simulate hurricane-level flooding of a coastal forest. Monitoring real-time soil conditions and tree physiological responses, we observed consistent impacts on soil biogeochemistry aligned with belowground drivers of tree mortality, but no consistent responses in aboveground vegetation. Our findings provide unprecedented empirically based insight into the earliest stages of a hypothesized forest mortality spiral and offer critical benchmarks for predicting coastal forest resilience in the face of accelerating climate change.

**Significance Statement:** Changing sea levels and storms are causing more flooding in coastal forests. This flooding kills trees, changing how coastal ecosystems function, but we do not fully understand what factors determine whether forests survive flooding. We designed an ecosystem-scale experiment to answer this question, with controlled saltwater and freshwater floods equivalent to a hurricane in experimental forest plots. Flooding quickly changed soil conditions, but we have not yet observed consistent tree stress responses. Our study provides the most detailed measurements to date of how coastal forests respond to flooding in real time. These findings will help us better understand the early mechanisms and warning signs of forests threatened by flooding.

## Introduction

Sea-level rise and shifting storm regimes are increasing the frequency and severity of flood events in low-lying coastal forests, driving their conversion to wetland or submerged ecosystems (1–5). This has dramatic consequences for carbon stocks (6), greenhouse gas emissions (7–9) and other biogeochemical or ecological phenomena that are fundamental to ecosystem structure and function. In addition, such landscape conversions have implications for coastal communities, as over 600 million people live in low-lying coastal areas worldwide (10, 11). Although it is well established that sea level rise leads to landscape transition and shifts in ecosystem states, the specific mechanisms through which flooding and seawater exposure initiate these changes trigger coastal forest mortality remain poorly understood (12).

Our understanding of the mechanisms and drivers of coastal forest mortality is particularly limited by the complex interactions between belowground and aboveground processes (13, 14). Changes in belowground conditions, including soil structure and biogeochemistry, impact aboveground woody plant health (12), and have been linked to formation of “ghost forests” (2, 15). Recent work suggests a combination of hypoxia and salinity drives mortality via hydraulic failure — blockage of water moving through xylem within a tree to the point of dehydration— followed by carbon starvation — insufficient carbohydrates to survive. The hypoxia and salinity thresholds required to instigate ecosystem conversion remain unknown, though previous literature suggests several days of persistent hypoxia is required to impact aboveground vegetation (16–18).

Historically, researchers have primarily gained mechanistic insights into the drivers and mechanisms of flooding disturbance on coastal forests through opportunistic observations, inferences from modeling studies, or space-for-time substitution studies (19–22). While valuable, these approaches typically focus on the later stages of ecosystem state change when stress in aboveground trees is visually apparent. This limitation stems from two challenges: the unpredictability of natural flooding events, and a frequent lack of concurrent belowground measurements. Controlled experimental manipulations therefore provide a crucial alternative to capture the earliest forest responses to flooding, particularly the initial belowground changes which theory predicts initiate the tree mortality process (12).

We established the manipulative, ecosystem-scale TEMPEST experiment to measure belowground and aboveground responses to coastal flooding over a decade to address this knowledge gap (23). In 2022, we flooded the TEMPEST plots with one event equivalent to Hurricane Sandy (15 cm), and in 2023, we flooded the plots with two consecutive events of the same magnitude. We monitored belowground and aboveground responses through a combination of in-situ soil (moisture, salinity, dissolved oxygen and redox) and tree stem sap flow sensors, greenhouse gas flux chambers, and manual measurements of greenhouse gas concentrations and leaf stress metrics to quantify responses of soil biogeochemistry and aboveground vegetation to flooding.

This study examines the 2023 flooding events as the first year with a comprehensive dataset to evaluate how short-term flooding can push coastal forests toward the tipping point between ecological stability and ghost forest formation. Using a hypothesized tree mortality spiral (12, 24) as a conceptual roadmap, we examine: 1) how soil and vegetation measurements related to changing biogeochemistry and tree mortality respond to flooding, and 2) whether these measurements provide evidence of either early (hydraulic failure-related) or advanced (carbon starvation-related) mortality. We hypothesize that 1) soils will respond rapidly to flooding conditions but return to pre-flood conditions quickly, while vegetation will not exhibit clear signs of stress, and 2) no evidence of carbon starvation and limited evidence of hydraulic failure will result from short-term flooding.

## Results

### Flooding drives soil saturation and electrical conductivity responses

We simulated two hurricane-scale events, flooding the Freshwater plot with municipal freshwater, and the Saltwater plot with brackish water from the nearby estuary (salinity ∼ 7 PSU) over two 10-hour flooding events starting 24 hours apart (Figure S1). No water was applied to the Control plot and no measurable precipitation occurred during these flooding treatments. We measured soil properties (volumetric water content, or VWC, and electrical conductivity, or EC) to track freshwater/brackish water delivery and soil salinity (Figure 1). Flooding increased VWC significantly in both treatment plots relative to the Control plot (p<0.05, Figure 1A, Table S1) by an average of 13% and 22% above pre-flood baseline in Saltwater and Freshwater plots, respectively (Table S2). VWC increased twice as fast during the first hour of flooding in the Freshwater plot compared to the Saltwater plot (14% and 7% above pre-flood baseline, respectively). Consistent with the conductivity of source water used to flood each treatment plot, soil EC increased significantly in both treatment plots relative to the Control plot (p<0.05, Figure 1B, Table S1), increasing 1,827% and 43% above baseline in Saltwater and Freshwater plots, respectively (Table S2). In the 5 days following the flood events, VWC returned to pre-flood values in the Saltwater plot but remained elevated in the Freshwater plot, and EC remained elevated by more than an order of magnitude in the Saltwater plot (Figure 1B), contradicting our our hypothesis of rapid soil recovery. VWC and EC both changed <1% in response to flooding in the Control plot (Figure 1, Table S2).

**Figure 1.**
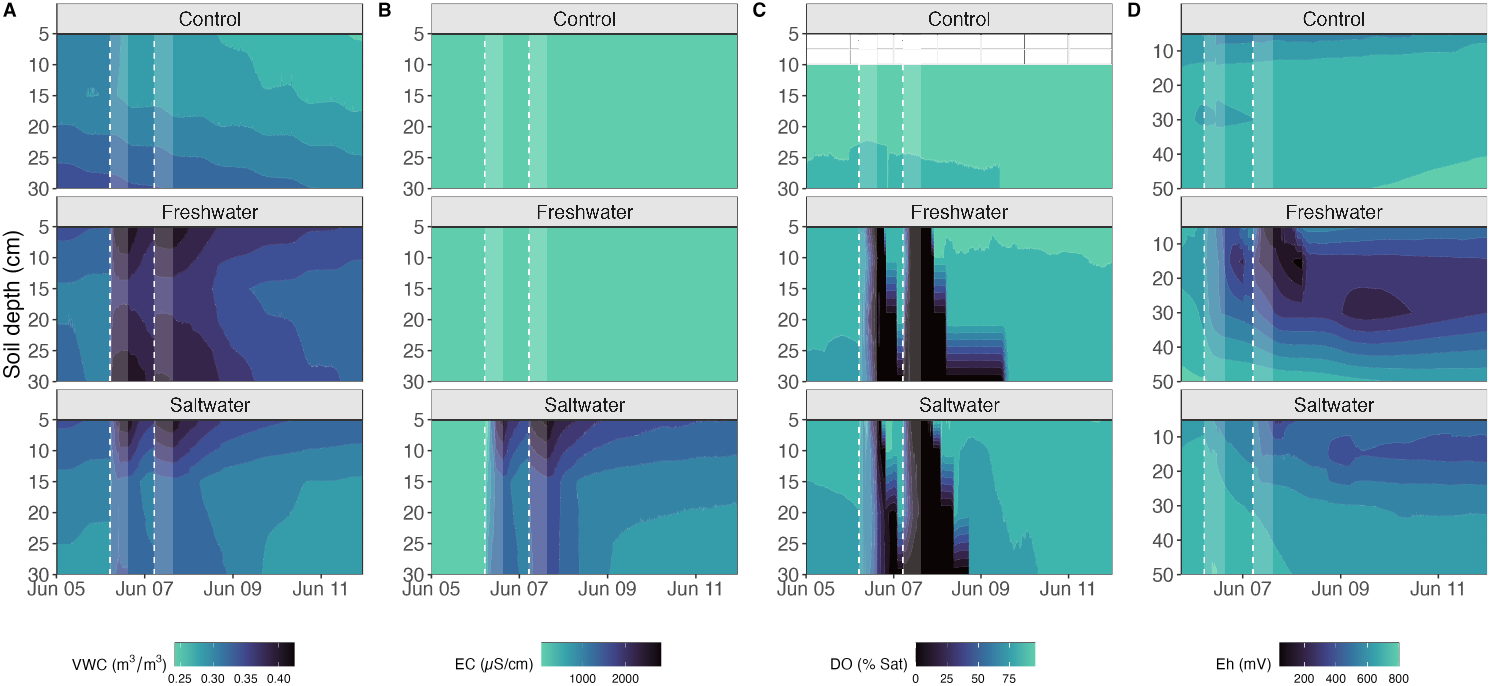
Depth-resolved time-series for soil measurements of A) volumetric water content (VWC), B) electrical conductivity (EC), C) dissolved oxygen (DO), and D) reduction-oxidation potential (Eh) measured in the Control and treatment (Saltwater and Freshwater) plots before, during, and after consecutive flooding events in June 2023. The start of each flood event is marked by a vertical dashed white line and shaded for the duration of flooding treatments. Higher VWC values indicate higher levels of soil saturation associated with flooding, and higher EC values indicate increased soil salinity. Lower DO values indicate increasingly limited oxygen availability, and lower Eh values indicate more reduced conditions.

### Flooding drives short-lived hypoxia but persistent reducing conditions

Prior to flooding, soil dissolved oxygen (DO) was equilibrated with the atmosphere, averaging 86-96% air saturation (Figure 1C, Table S2). During flooding, DO dropped significantly in both treatment plots relative to the Control plot (p<0.05, Figure 1C, Table S2), reaching hypoxia (insufficient oxygen availability, defined as <21% air saturation, see Methods) with minima <1%. DO decreased ∼13% in the Control plot over the course of the study (Table S2) but showed no response to flooding (Figure 1C). Following the first flooding event, soil DO recovered quickly in the top 15 cm but remained hypoxic in both treatment plots at 30 cm (Figure 1C). After the second flooding event, all depths returned to the pre-flood range (80-90% air saturation) within 3 days (Figure 1C, Figure S2). We quantified oxygen consumption rates across treatment plots, which averaged 11% air saturation per hour in the Freshwater plot and 7% air saturation per hour in the Saltwater plot during the first 2023 flooding event (Figure S2). On average across treatment plots, oxygen consumption rates were faster during the second flooding event (12% air saturation per hour) relative to the first flooding event (average 7% air saturation per hour). Hypoxia persisted longer with increasing depth (2-3 days at 30 cm compared to ∼1.4 days at 5 cm), and longer in the Freshwater plot (1.4-3 days) than the Saltwater plot (1.3-2 days). In both treatment plots, all depths reached values <1% air saturation (anoxia) during the second flooding event, persisting less than 7 hours to almost 3 days depending on depth.

Consistent with DO responses, reduction-oxidation (redox) potential relative to the standard hydrogen electrode (Eh) decreased significantly during flooding in both Freshwater and Saltwater plots relative to the Control plot (p<0.05, Figure 1D, Table S1). During flooding, Eh decreased 26% and 10% on average in Freshwater and Saltwater plots, respectively, but increased 3% in the Control plot (Figure 1D, Table S2). In contrast to DO, and contrary to our hypothesis of rapid soil recovery, Eh values had not returned to pre-flood values 5 days after flooding treatments stopped (Figure 1D, Table S2). We observed a change from oxic to weakly reducing conditions (200-400 mV) at 5 cm but no deeper for the Saltwater plot, persisting for less than 6% of the study period (Figure S3). In the Freshwater plot, weakly reducing conditions across 5-30 cm persisted for up to 87% of the study period, with moderately reducing conditions (−100 to 200 mV) at 5 and 15 cm lasting ∼10% of the period (Figure S3).

### Flooding alters CO_2_ dynamics in soils but not tree stems

We explored flooding responses in soils and tree stems through changes in concentrations and fluxes of CO_2_, a volatile and important component of the terrestrial carbon cycle, to see if the changes in soil biogeochemistry propagated to aboveground vegetation (Figure 2). Prior to flooding, soil CO_2_ fluxes were not significantly different between any of the plots (Figure 2A). During flooding, soil CO_2_ fluxes were significantly lower (p<0.05) in the treatment plots (by 52% and 60% in the Freshwater and Saltwater plots, respectively) and significantly higher (by 70%) in the Control plot. Following flooding, soil CO_2_ fluxes returned to pre-flood values the day after treatments stopped. Soil CO_2_ concentrations showed similar patterns to soil CO_2_ fluxes, where treatment plot values decreased during flooding and returned to pre-flood levels following flooding, although the Control plot did not show an equivalent increase in soil CO_2_ concentrations during the events (Figure 2C). In contrast, tree stem CO_2_ fluxes and concentrations did not show significant responses to flooding, except for an increase during flooding in Saltwater tree stem CO_2_ concentrations (Figure 2B and D). Together, soil and tree stem CO_2_ responses (or lack thereof) to flooding support our first hypothesis, where clear changes in soil biogeochemistry did not propagate to aboveground vegetation.

**Figure 2.**
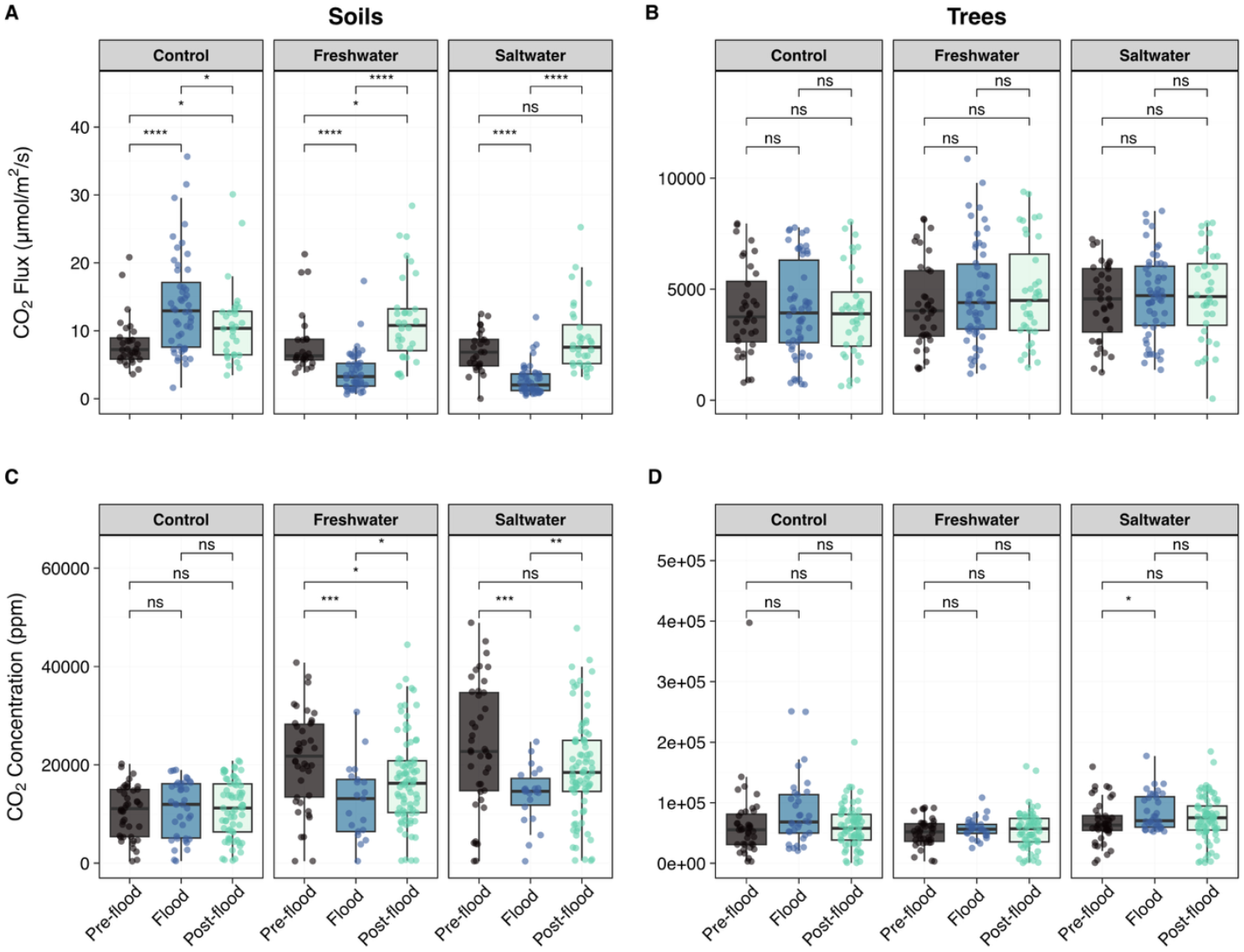
Measurements of soil carbon dioxide (CO2) A) fluxes and B) concentrations made prior to (“Pre-flood”), during (“Flooded”), and after (“Post-flood”) sequential flooding events. Equivalent measurements are presented for tree stem CO_2_ fluxes (C) and concentrations (D). Significance pairwise differences are shown for as ns (p>0.05), * (p<0.05), ** (p<0.01) or *** (p<0.001).

### Flooding drives inconsistent vegetation responses

To quantify vegetative stress in the canopy, we measured two leaf-level physiology metrics, photosynthesis rates and stomatal conductance, which we expect to decrease in response to flooding (12). However, we observed inconsistent and somewhat conflicting indications of stress, partially contradicting our hypothesis that vegetation would not exhibit clear stress signals (Figure S4). Stomatal conductance significantly decreased (p<0.05) in both treatment plots, agreeing with our expectations of flood-induced stress (Figure S4A). Unexpectedly, photosynthesis rates, which we expected to decrease in response to stress, increased significantly in the Freshwater plot, and did not change in the Saltwater plot (Figure S4B).

Stress via hydraulic failure in response to flooding is expected to decrease sap flux in tree stems. Since this response may be lagged and data were available, we analyzed the entire 2023 growing season for the three dominant tree species (American beech, red maple, and tulip poplar) in our plots (Figure 3). We observed differences between Control and treatment plots during the growing season (Figure 3A), where sap flux was generally higher in the Freshwater plot relative to Control, while sap flux was generally lower in the Saltwater plot relative to Control, and tulip poplars in the Saltwater exhibited earlier senescence. Normalized sap flux trends increased in beech trees in both plots, but decreased in red maples and tulip poplars (Figure 3B. We found significant post-flood Sen-Theil slopes (Table S3) for tulip poplar in Freshwater and all three species in Saltwater (Table S3). Tulip poplars in Saltwater showed the steepest trend, with a slope twice as steep as tulip poplars in Freshwater, matching observations of earlier senescence (Figure 3A).

**Figure 3.**
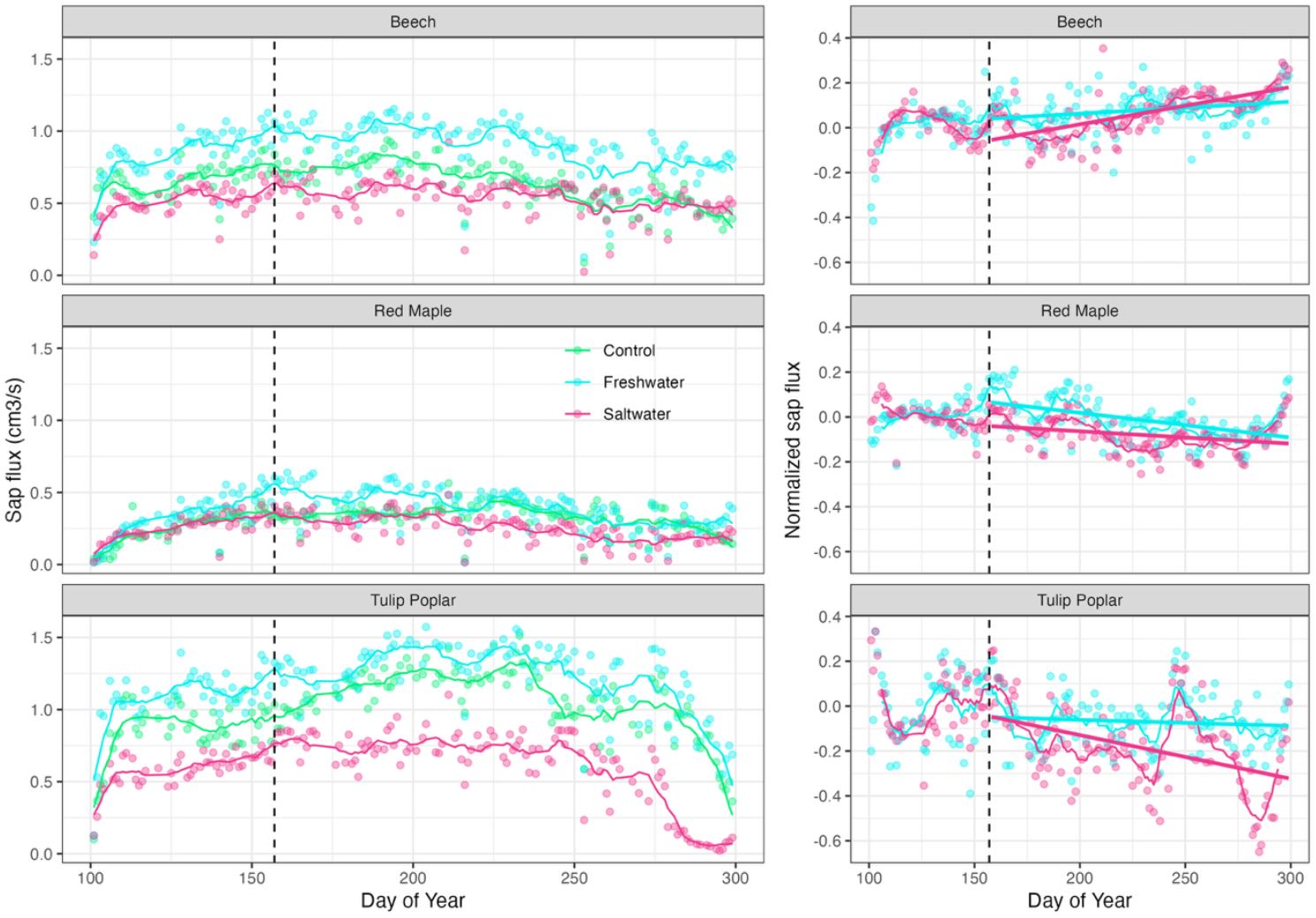
Tree stem sap flux responses by plot and species across the 2023 growing season to flooding. A) time-series of sap fluxes with the start of flooding marked by a vertical dashed line, and B) sap fluxes normalized to pre-treatment values and the Control plot following McDowell et al. (50) with linear regression lines fit to post-flood data. Sen-Theil slopes and their significance are presented in Table S3

## Discussion

### Short-term flooding drove dramatic changes in soil biogeochemistry

Consistent with our hypothesis, soil biogeochemistry rapidly responded to short-term flooding across several hypothesized drivers of coastal tree mortality, including soil hypoxia and salinity (12, 18). Changes in DO support our hypothesis and are consistent with rapid consumption and re-equilibration of soil oxygen in response to flooding observed by others (25, 26). Hypoxia observed across the top 30 cm of the soil profile (Figure 1C) is a well-documented response to flooding in forest soils (23, 27–29), and a hypothesized key driver of ghost forest formation (12). Our observations are consistent with increasing soil moisture reducing oxygen diffusion rates below levels required to meet biological and chemical oxygen demand, thereby driving hypoxia (30, 31). Previous studies quantifying the duration of hypoxia needed to alter tree function vary widely in their methods (including age and species of tree, and environment), but provide a broad envelope ranging from as little as 3 to 5 days (16) to several weeks (17) of persistent hypoxia to impact seedlings. We also note that species and tree age play important roles in flood tolerance (32), and quantitative hypoxia studies are primarily conducted on seedlings, while our forest plots contain mature trees. We thus would not expect a duration of < 4 days to inflict measurable hypoxia-driven damage on the trees in our treatment plots.

The rates of oxygen consumption during the first flood event (declining to < 1% in 9-10 hours) are consistent with rates reported in laboratory experiments (9-18 hours) (25). Interestingly, when we compared oxygen consumption rates between flooding events, all sites that returned to an oxic state between the first and second flood event exhibited faster oxygen consumption rates during the second flood event (117-221% of rates observed during the first flooding event). We attribute this difference in oxygen consumption to faster flood water infiltration rates that would be expected from the higher antecedent soil moisture prior to the second flooding event. Respiration is typically most strongly impacted during the first drying/rewetting cycle in soils (33), supporting our explanation that diffusion rather than biological or chemical processes controlled oxygen consumption. Additionally, we saw faster increases in VWC during the first flood in Freshwater compared to Saltwater, which matches faster rates of DO consumption and longer hypoxia in the Freshwater plot, potentially due to more rapid declines in oxygen diffusion rates.

In contrast to DO and our hypothesis, redox decreased more slowly and did not recover to pre-flood values during the study period (Figure 1D). The temporal decoupling of oxygen and redox dynamics is consistent with other studies, including flooding across coastal terrestrial-aquatic interfaces (34), attributed to redox poising capacity (35). Persistent reducing conditions with lower Eh values in the Freshwater plot relative to the Saltwater plot matches our observations of higher VWC, more rapid increasing VWC, more rapid DO consumption, and more persistent hypoxia in the Freshwater plot. Persistent, more reducing conditions in the Freshwater plot also match higher VWC and more persistent hypoxia in Freshwater compared to the Saltwater plot, which is potentially related saltwater-induced changes in soil permeability (36). These persistent reducing conditions may be precursors to longer-term soil biogeochemical changes that could eventually impact root function.

The impacts of short-term flooding on soil biogeochemistry also included a temporary reduction in soil CO_2_ fluxes (Figure 2), suggesting either a change in source or transport. Because concentrations and fluxes of CO_2_ are measured via two different methods, at different times and in different locations within the plots, matching flux and concentration patterns suggest that changes in fluxes are explained by a change in concentration along with an expected decrease in diffusion rates with increased VWC (Figure 1A). Other studies of forest soils report both increases and decreases in CO_2_ fluxes in response to flooding (8, 37, 38). Flooding can drive rapid increases in soil respiration via several mechanisms, including a sudden increase upon rewetting known as the “Birch effect” (39), and degassing of air-filled pores (40). However, we observed the opposite response, suggesting that other mechanisms are more likely, including oxygen limitation slowing aerobic respiration (41).

### Short-term flooding did not drive consistent vegetation impacts

We did not observe consistent responses of aboveground vegetation to our short-term flooding events (Figure 3), which agrees with our first hypothesis and suggests the limited duration of flooding was not sufficient to manifest vegetation disturbance signatures. This is supported by the lack of response in tree stem CO_2_ concentrations and emissions (Figure 2B/D), which we expected to follow soil CO_2_ patterns (Figure 2A/C). Inconsistent responses between stomatal conductance (which matched our expectation, decreasing in both treatment plots, Figure S4A) and photosynthetic rates (which differed in flooding responses between treatment plots) further suggest that clear flood-induced changes to soil oxygen, redox, and CO_2_ did not consistently impact aboveground vegetation. Our experiments indicate that tree death from flooding takes more than one event, even when the belowground drivers change as expected.

The temporal extent of hypoxia required to initiate events hypothesized to impact tree hydraulics and carbon allocation in coastal forests remains unknown. Our experiment was conducted during a relatively dry period, where the benefit of increased soil saturation may have partially counteracted the negative impacts of hypoxia. Since vegetation responses can lag behind disturbances (42), our full growing season of sap flux data provides a valuable initial metric, indicating tulip poplars in the Saltwater plot exhibit both reduced sap flux and a shortened growing season. This may partially refute our hypothesis by presenting initial evidence that hydraulic failure may be occurring but also indicates species-specific and salinity-specific responses. The strongest responses observed consistently in the Saltwater plot suggest that salinity-driven changes may be as significant, if not more so, than hypoxia-driven changes in initiating hydraulic failure. If future data verify that this period signifies the start an irreversible ecological change in the Saltwater plot, our results suggest that tulip poplars could serve as an effective sentinel species (43) for future flood-induced forest disturbances in eastern North America.

### Short-term flooding impacts on coastal tree mortality

We used our measurements of soil and vegetation responses to flooding disturbances to assess how far the 2023 flooding events pushed our treatment plots along the hypothetical tree mortality spiral (Figure 4). Since the TEMPEST experiment is focused on flooding as the primary disturbance, several mechanisms (storm damage, drought, elevated temperature, elevated VPD, and elevated CO_2_) are outside the scope of this study and cannot be assessed. However, our diverse datasets enable us to evaluate how far into the spiral impacts were observed, enhancing our understanding of how short-term flooding can drive a forest towards the tipping point of tree mortality and ghost forest formation through two hypothesized mortality mechanisms: hydraulic failure and carbon starvation.

**Figure 4.**
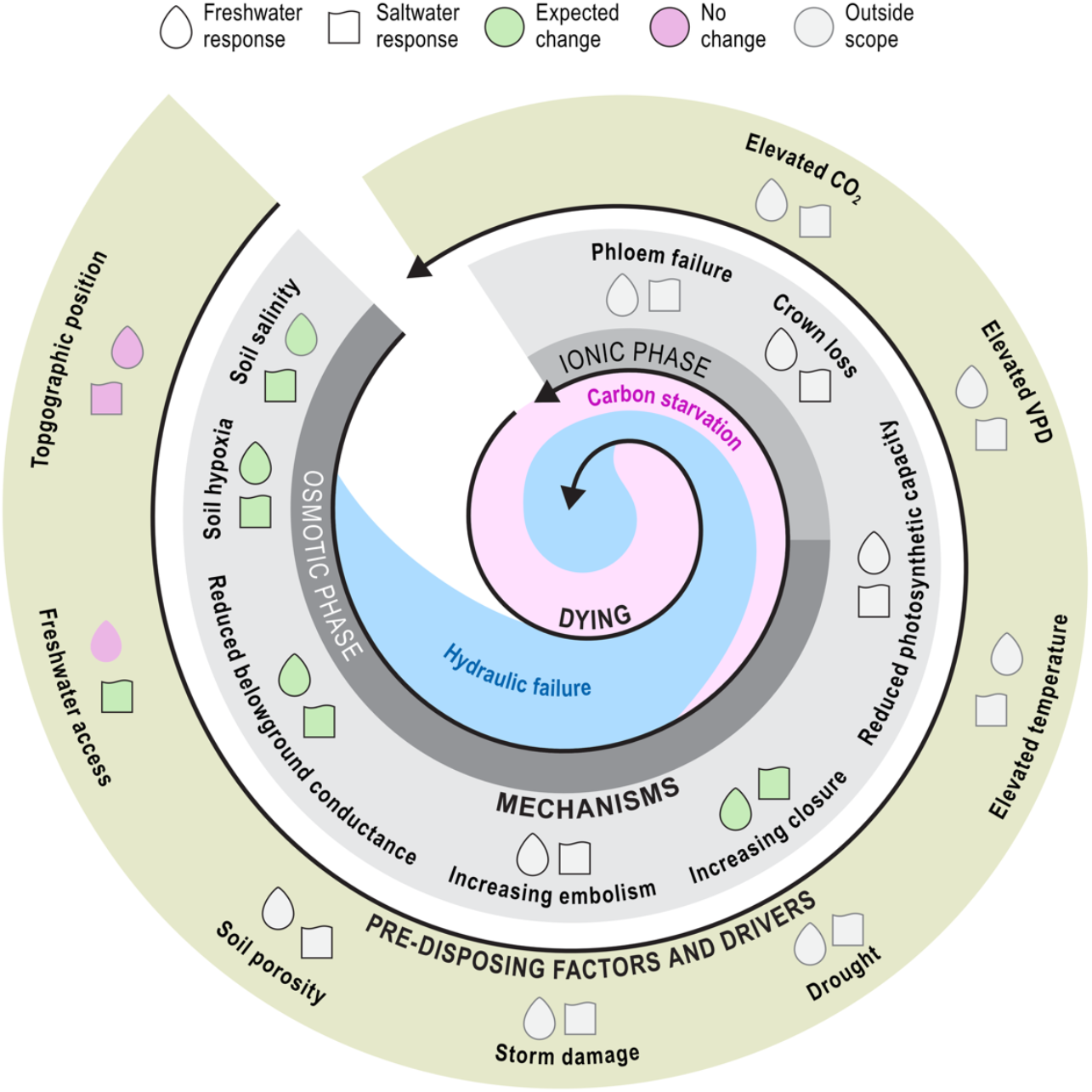
Conceptual figure adapted from McDowell et al. (12), showing hypothesized drivers and mechanisms of tree mortality spiral, and empirical results based on our datasets.

We measured changes in several belowground drivers to explore hypothesized mechanisms such as soil salinity (Saltwater plot only, Figure 1B), freshwater access (Freshwater plot only, Figure 1A), and soil hypoxia (both treatment plots, Figure 1C). We inferred belowground hydraulic conductance from sap flux and water potentials, which, contrary to our conceptual understanding, was higher (though not significantly so) in both treatment plots (Figure S5). Although neither redox nor soil respiration impacts are explicitly included in the original framework, both are strongly linked to soil saturation and associated hypoxia, so we suggest both factors should be located between soil hypoxia and reduced conductance, as additional markers of progress down the mortality spiral. Soil drivers of tree mortality thus consistently indicate conditions primed for onset of hydraulic failure.

We observed consistent stress responses through stomatal closure in both treatment plots (Figure S4B), yet photosynthesis unexpectedly increased, likely due to increased freshwater access to trees in the Freshwater plot (Figure 1A, S4A). Although this may reflect greater photosynthetic capacity, our single-time-point measurement does not provide enough information to interpret this trend confidently, so we left the photosynthesis metric blank in Figure 4. Mechanistically, stomatal behavior, photosynthesis, and sap flow are closely linked: maintaining carbon gain while minimizing water loss is essential, especially under saline conditions where plants must also manage salt uptake. Salt stress can trigger stomatal closure to reduce water loss, but this also limits CO_2_ uptake, creating a physiological trade-off that influences photosynthesis and overall plant function. Balancing carbon gain and water use in the context of salt uptake is critical for protecting metabolic pathways and maintaining leaf longevity (44).

Together, the responses in Figure 4 predict a stronger impact on the Saltwater plot (increased soil salinity) relative to the Freshwater plot. The species-specific nature of sap flux responses may indicate varying thresholds of flood tolerance, with tulip poplars appearing most sensitive, red maples showing moderate responses, and beech trees demonstrating resilience through increased sap flux. Synthesis of our findings as they relate to the tree mortality spiral therefore suggests that the flooding regime our treatment plots have been subjected to thus far are 1) not enough to notably impact the forest in the Freshwater plot, and 2) may already be pushing the Saltwater plot across an ecological tipping point. The sustained soil EC changes, decreased stomatal conductance, and declining sap flux in salt-sensitive species together suggest the initial stages of a transition that could accelerate with subsequent flooding events. Notably, the Saltwater plot tulip poplar trees experienced an early senescence event in 2024 (personal observation, not reported further here), supporting this inference that the disturbance appears to be related to salinity rather than solely flooding. While our early observations are not enough to substantiate vegetation impacts via hydraulic failure, continued monitoring will inform if we are capturing a tipping point or if these are temporary impacts on a resilient forest.

As coastal forests experience increasing flooding pressure in tandem with other stressors (5, 12, 45), it is essential to understand how the extent, intensity, and duration of inundation events alter forest health, including soil biogeochemistry and tree physiology. Because of the intrinsic links between belowground and aboveground forest processes (13, 14), collecting simultaneous belowground and aboveground datasets during flooding disturbance is critical for improving process representation of coastal forest vulnerability to flooding in Earth system models, and how their responses impact coastal carbon cycling (46). While our data snapshot suggests our experimental plots may be at an ecological tipping point between healthy and dying forest (Figure 4), our analyses also highlight the complexity of forest responses to flooding, and the value of ecosystem-scale experiments to provide mechanistic insights. Our findings demonstrate that even short-term flooding events may initiate the early stages of the tree mortality spiral when brackish or saline water is involved, highlighting the vulnerability of coastal forests to storm surge and sea level rise rather than increased precipitation alone. Sustained, longer-term observations will be needed to identify belowground and aboveground signatures predicting future ecosystem transitions, providing much-needed quantitative tools for managing coastal forest resources.

## Materials and Methods

### Site Description

The Terrestrial Ecosystem Manipulation to Probe the Effects of Storm Treatments (TEMPEST) experiment is located in Edgewater, MD, USA at the Smithsonian Environmental Research Center’s Global Change Research Wetland. In 2019-2020, three 40m x 50m plots were established in the upland coastal forest adjacent to a tidal marsh: Freshwater (a treatment plot to be flooded with freshwater), Saltwater (a treatment plot to be flooded with estuarine water), and Control (a control plot not subjected to any flooding treatment). All plots were outfitted with a variety of sensors to monitor groundwater, soils, and vegetation, described in more detail below. Site lithology features approximately half a meter of organic-rich soils overlaying an approximately meter-thick silty clay layer perched on at least 1.5 meters of sandy soils (47). A full description of the TEMPEST experiment can be found in Hopple et al. (23).

### Flooding Events

TEMPEST is designed to accelerate flooding pressure by increasing the number of flooding events each year. In 2022, the treatment plots were flooded with a single event. In 2023 (the period for this study), the treatment plots were flooded with two flood events. The first 2023 flooding event started at 6 June 2023 05:00 EST and the second started 24 hours later. Both events lasted 10 hours, with each event delivering ∼80,000 gallons (∼303,000 L) of water to each treatment plot, totaling 360,000 gallons (∼1.36 million L) applied to the treatment plots over both events, or ∼15 cm of water across each plot. Water applied to the Freshwater plot was sourced from a municipal water source, and water applied to the Saltwater plot was pumped from the adjacent Rhode River estuary (∼ 7 PSU), first passing through a filter to remove debris >125 µm.

### Measurements

All measurements used in this manuscript are summarized in Table S4. These measurements included sensors deployed in the soil and in tree stems, and samples collected from the soil, from tree stems, and from the tree canopy. The sections below provide details on each of these measurements.

### Soil sensors

For soil sensor datasets, we used data from 5-12 June, 2023 to make our visual and statistical comparisons as consistent as possible across sensors. We measured soil electrical conductivity (EC) and volumetric water content (VWC) using TEROS 12 sensors deployed at 5, 15, and 30 cm in multiple grid-cells across each plot. We measured dissolved oxygen (DO) using Firesting optical sensors (Pyroscience), using a 4-channel logger with 4 DO sensors and one temperature sensor for temperature compensation at 5, 10, 20, and 30 cm depths in both treatment plots, and using three handheld Firesting GO sensors at 10, 20, and 30 cm depths in the Control plot. All DO sensors were calibrated at the same time in the same environment to percent air saturation immediately prior to deployment. Firesting 4-channel systems are designed for laboratory conditions, and we developed a custom battery-powered, weather-resistant setup to enable continuous logging for the experiment’s duration. We defined hypoxia based on Armstrong et al. (48) as 4.5 kPa, which equals approximately 21% air saturation.

We measured redox (reduction-oxidation potential) using 4-electrode redox probes (SWAP) with depths of 5, 15, 30, and 50 cm. In each plot, 5 identical redox sensors were deployed in a 1 m diameter circle, and two reference electrodes were deployed on either side of the circle. Each redox sensor nest was powered and logged using an CR1000x logger (Campbell Scientific) attached to an isolated power source to avoid electrical interference.

### Greenhouse gas measurements

We measured soil CO_2_ fluxes from four 20-cm PVC collars permanently installed in random configurations in each treatment plot, with ∼8 cm of the color above the soil and ∼2 cm below. Fluxes were measured using a LI-COR 8200-01S smart chamber connected to an LI-COR 7810 infrared gas analyzer in two 60-second replicates 2-4 times per day in all treatments prior to, during, and after the flooding events. CO_2_ fluxes from tree stems were also measured using a LI-COR 7810. Flux collars were sealed to stems with silicone and interfaced to the gas analyzer with plates attached to the collars with elastic cord. Soil CO_2_ concentrations were collected from five gas well nests located across each treatment plot (co-located in grid cells with TEROS, Firesting, and SWAP sensors) in exetainers evacuated under N_2_. For gas well samples containing water, we extracted CO_2_ from water using a 2:1 dilution of air to water and 2 minutes of shaking to equilibrate prior to transferring 20 mL of gas to the exetainer. Air reference samples were collected at all plots for each sampling point to correct for atmospheric concentrations. Tree stem CO_2_ concentrations were collected and analyzed in the same way as soil CO_2_ concentrations but from trees with stainless steel pipes installed in bore holes at breast height; pipes were sealed with gas-tight septa (49).

### Sap flow sensors

We measured sap flux using sap flow sensors following the thermal dissipation method where two temperature probes (Plant Sensors) with a 2-cm sensing length are installed 10 cm apart, with the top probe heated at constant voltage and the bottom probe functioning as a reference. Sensors were installed in 6 individuals for each of the three dominant tree species (*Fagus gradifolia* or American beech, *Acer rubrem* or red maple, and *Liriodendron tulipifera* or tulip poplar), totaling 18 trees across each plot. We followed published methods to calculate sap flux densities from voltages (29). We normalized sap flux densities following McDowell et al. (50) to assess change in sap flux relative to 1) pre-flood conditions within each plot and species, and 2) relative to the Control plot.

### Leaf measurements

We made manual measurements of photosynthesis, water potential, and stomatal conductance on leaves collected 5 days after flooding. Intercellular data was measured on freshly collected leaf samples for which there are practical constraints due to height to the first branch for many focal trees. Photosynthesis and stomatal conductance were measured on leaves collected on all American beech and some red maples with sap flow sensors. We used a LI-6400/XT to measure both intercellular variables and a pressure chamber to measure pre-dawn and midday water potentials. We estimated belowground conductance as the quotient of sap flux density and the difference between pre-dawn and midday water potentials.

### Statistics

DO consumption rates were calculated as the difference between starting DO and minimum DO concentrations during each flooding event, except for 30 cm, where oxygen remained low following the first flooding event, resulting in no rates for the second flooding event at 30 cm. We calculated Eh (redox potential) following Machado-Silva et al. (34) to report our measurements relative to a standard reference and calculated redox categories as: oxidizing (Eh ≥ 400 mV), weakly reducing (200 mV ≤ Eh < 400 mV), moderately reducing (−100 mV ≤ Eh < 200 mV), and strongly reducing (Eh < −100 mV).

For statistical comparisons of significant (p<0.05) differences between groups, we first tested each dataset for normality using Shapiro-Wilk’s tests and homogeneity of variance using Levene’s tests. For time-series collected by soil sensors, we visually examined autocorrelation functions to test the assumption of independence. Because all soil sensor datasets in Table S1 exhibited non-normal distributions, different variances between groups and strong autocorrelation, we used non-parametric repeated measures Friedman tests to compare between treatment plots and control plots separately for pre-flood, flood, and post-flood time-periods. Based on the results of these tests, we then used post-hoc pairwise Wilcoxon tests to compare between each pair of treatment plots for each time-period. For data that did not violate the assumption of independence but violated either the assumption of normality or homogeneity of variance, we used Wilcoxon tests to compare between each treatment. All analyses were conducted in R v4.3 using code available on Github (https://github.com/COMPASS-DOE/tempest2_do).

## Acknowledgments

We thank the field sampling and infrastructure teams that make the TEMPEST experiment possible, particularly Anya Hopple, Evan Phillips and Leticia Sandoval for their contributions that made this work possible. We also thank James Stegen for his early conceptual contributions to TEMPEST’s experimental design, and Nathan Johnson for graphical editing of the conceptual figure. This research was supported by COMPASS-FME, a multi-institutional project supported by the U.S. Department of Energy, Office of Science, Biological and Environmental Research as part of the Environmental System Science Program, and by the Smithsonian Environmental Research Center. The Pacific Northwest National Laboratory is operated for DOE by Battelle Memorial Institute under contract DE-AC05-76RL01830.

## References

1. Y. Chen, M. L. Kirwan, A phenology- and trend-based approach for accurate mapping of sea-level driven coastal forest retreat. Remote Sensing of Environment 281, 113229 (2022).

2. M. L. Kirwan, K. B. Gedan, Sea-level driven land conversion and the formation of ghost forests. Nat. Clim. Chang. 9, 450–457 (2019).

3. N. W. Schieder, M. L. Kirwan, Sea-level driven acceleration in coastal forest retreat. Geology 47, 1151–1155 (2019).

4. J. Z. Sippo, C. E. Lovelock, I. R. Santos, C. J. Sanders, D. T. Maher, Mangrove mortality in a changing climate: An overview. Estuarine, Coastal and Shelf Science 215, 241–249 (2018).

5. J. B. Cannon, C. J. Peterson, C. M. Godfrey, A. W. Whelan, Hurricane wind regimes for forests of North America. Proceedings of the National Academy of Sciences 120, e2309076120 (2023).

6. L. S. Smart, et al., Aboveground carbon loss associated with the spread of ghost forests as sea levels rise. Environ. Res. Lett. 15, 104028 (2020).

7. J. Batson, et al., Soil greenhouse gas emissions and carbon budgeting in a short-hydroperiod floodplain wetland. Journal of Geophysical Research: Biogeosciences 120, 77–95 (2015).

8. S. Petrakis, A. Seyfferth, J. Kan, S. Inamdar, R. Vargas, Influence of experimental extreme water pulses on greenhouse gas emissions from soils. Biogeochemistry 133, 147–164 (2017).

9. H. Chen, et al., Unique biogeochemical characteristics in coastal ghost forests – The transition from freshwater forested wetland to salt marsh under the influences of sea level rise. Soil & Environmental Health 1, 100005 (2023).

10. B. Neumann, A. T. Vafeidis, J. Zimmermann, R. J. Nicholls, Future Coastal Population Growth and Exposure to Sea-Level Rise and Coastal Flooding - A Global Assessment. PLOS ONE 10, e0118571 (2015).

11. J. P. Hochard, S. Hamilton, E. B. Barbier, Mangroves shelter coastal economic activity from cyclones. Proc. Natl. Acad. Sci. U. S. A. 116, 12232–12237 (2019).

12. N. G. McDowell, et al., Processes and mechanisms of coastal woody-plant mortality. Global Change Biology 28, 5881–5900 (2022).

13. Y. Preisler, F. Tatarinov, J. M. Grünzweig, D. Yakir, Seeking the “point of no return” in the sequence of events leading to mortality of mature trees. Plant, Cell & Environment 44, 1315–1328 (2021).

14. W. M. Hammond, et al., Global field observations of tree die-off reveal hotter-drought fingerprint for Earth’s forests. Nat Commun 13, 1761 (2022).

15. K. W. Krauss, et al., Ghost forests of Marco Island: Mangrove mortality driven by belowground soil structural shifts during tidal hydrologic alteration. Estuarine, Coastal and Shelf Science 212, 51–62 (2018).

16. H. Folzer, J. F. Dat, N. Capelli, D. Rieffel, P.-M. Badot, Response of sessile oak seedlings (Quercus petraea) to flooding: an integrated study. Tree Physiol 26, 759–766 (2006).

17. C. Parent, M. Crèvecoeur, N. Capelli, J. F. Dat, Contrasting growth and adaptive responses of two oak species to flooding stress: role of non-symbiotic haemoglobin. Plant, Cell & Environment 34, 1113–1126 (2011).

18. W. H. Conner, K. W. McLeod, J. K. McCarron, Survival and Growth of Seedlings of Four Bottomland Oak Species in Response to Increases in Flooding and Salinity. Forest Science 44, 618–624 (1998).

19. E. Worley, et al., Impacts of Hurricane Michael on Watershed Hydrology: A Case Study in the Southeastern United States. Forests 13, 904 (2022).

20. H. Niwa, M. Kamada, S. Morisada, M. Ogawa, Assessing the impact of storm surge flooding on coastal pine forests using a vegetation index. Landscape Ecol Eng 19, 151–159 (2023).

21. A. J. Smith, et al., Litter Decomposition in Retreating Coastal Forests. Estuaries and Coasts 47, 1139–1149 (2024).

22. J. Ding, et al., Modeling the mechanisms of coastal vegetation dynamics and ecosystem responses to changing water levels. EGUsphere 1–31 (2025). 10.5194/egusphere-2025-1544.

23. A. M. Hopple, et al., Attaining freshwater and estuarine-water soil saturation in an ecosystem-scale coastal flooding experiment. Environ Monit Assess 195, 425 (2023).

24. J. A. Duberstein, et al., Small gradients in salinity have large effects on stand water use in freshwater wetland forests. Forest Ecology and Management 473, 118308 (2020).

25. K. F. Patel, et al., Time to anoxia: Observations and predictions of oxygen drawdown following coastal flood events. Geoderma 444, 116854 (2024).

26. P. Regier, et al., Seasonal drivers of dissolved oxygen across a tidal creek–marsh interface revealed by machine learning. Limnology and Oceanography (2023). 10.1002/lno.12426.

27. J. P. Megonigal, W. H. Patrick Jr., S. P. Faulkner, Wetland Identification in Seasonally Flooded Forest Soils: Soil Morphology and Redox Dynamics. Soil Science Society of America Journal 57, 140–149 (1993).

28. P. Parolin, F. Wittmann, Struggle in the flood: tree responses to flooding stress in four tropical floodplain systems. AoB Plants 2010, plq003 (2010).

29. A. N. Myers-Pigg, et al., Short-term coastal forest responses to a hurricane-scale freshwater and saltwater flooding experiment. [Preprint] (2025). Available at: https://www.biorxiv.org/content/10.1101/2025.04.13.648622v1 [Accessed 6 May 2025].

30. Z. Fischer, P. Blažka, L. Dubis, Respiration Rates of Organic Soil Depending on Changes of Moisture and Aeration. Open Journal of Soil Science 7, 101–110 (2017).

31. Z. Fischer, L. Dubis, Soil Respiration in the Profiles of Forest Soils in Inland Dunes. Open Journal of Soil Science 9, 75–90 (2019).

32. N. Leksungnoen, W. Eiadthong, R. Kjelgren, Thailand’s catastrophic flood: Bangkok tree mortality as a function of taxa, habitat, and tree size. Urban Forestry & Urban Greening 22, 111–119 (2017).

33. E. A. de Nijs, L. C. Hicks, A. Leizeaga, A. Tietema, J. Rousk, Soil microbial moisture dependences and responses to drying–rewetting: The legacy of 18 years drought. Global Change Biology 25, 1005–1015 (2019).

34. F. Machado-Silva, et al., Short-Term Groundwater Level Fluctuations Drive Subsurface Redox Variability. Environ. Sci. Technol. 58, 14687–14697 (2024).

35. A. J. Burgin, T. D. Loecke, The biogeochemical redox paradox: how can we make a foundational concept more predictive of biogeochemical state changes? Biogeochemistry 164, 349–370 (2023).

36. D. Liu, M. Edraki, P. Fawell, L. Berry, Improved water recovery: A review of clay-rich tailings and saline water interactions. Powder Technology 364, 604–621 (2020).

37. E. M. Docherty, A. D. Thomas, Larger floods reduce soil CO2 efflux during the post-flooding phase in seasonally-flooded forests of Western Amazonia. Pedosphere 31, 342–352 (2021).

38. A. M. Hopple, S. C. Pennington, J. P. Megonigal, V. Bailey, B. Bond-Lamberty, Disturbance legacies regulate coastal forest soil stability to changing salinity and inundation: A soil transplant experiment. Soil Biology and Biochemistry 169, 108675 (2022).

39. H. F. Birch, The effect of soil drying on humus decomposition and nitrogen availability. Plant Soil 10, 9–31 (1958).

40. J. Muhr, W. Borken, Delayed recovery of soil respiration after wetting of dry soil further reduces C losses from a Norway spruce forest soil. Journal of Geophysical Research: Biogeosciences 114 (2009).

41. A. M. Hopple, S. C. Pennington, J. P. Megonigal, V. Bailey, B. Bond-Lamberty, Root and Microbial Soil CO2 and CH4 Fluxes Respond Differently to Seasonal and Episodic Environmental Changes in a Temperate Forest. Journal of Geophysical Research: Biogeosciences 128, e2022JG007233 (2023).

42. S. A. Kannenberg, C. R. Schwalm, W. R. L. Anderegg, Ghosts of the past: how drought legacy effects shape forest functioning and carbon cycling. Ecology Letters 23, 891–901 (2020).

43. M. Ardón, K. M. Potter, E. W. Jr, C. W. Woodall, Coastal carbon sentinels: A decade of forest change along the eastern shore of the US signals complex climate change dynamics. PLOS Climate 4, e0000444 (2025).

44. M. C. Ball, R. Munns, Plant Responses to Salinity Under Elevated Atmospheric Concentrations of CO2. Aust. J. Bot. 40, 515–525 (1992).

45. Y. Xie, X. Wang, J. A. Silander, Deciduous forest responses to temperature, precipitation, and drought imply complex climate change impacts. Proc. Natl. Acad. Sci. U. S. A. 112, 13585–13590 (2015).

46. N. D. Ward, et al., Representing the function and sensitivity of coastal interfaces in Earth system models. Nat Commun 11, 2458 (2020).

47. M. B. Adebayo, et al., A hydrogeophysical framework to assess infiltration during a simulated ecosystem-scale flooding experiment. Journal of Hydrology 626, 130243 (2023).

48. W. Armstrong, T. Webb, M. Darwent, P. M. Beckett, Measuring and interpreting respiratory critical oxygen pressures in roots. Ann Bot 103, 281–293 (2009).

49. N. D. Ward, et al., Longitudinal Gradients in Tree Stem Greenhouse Gas Concentrations Across Six Pacific Northwest Coastal Forests. Journal of Geophysical Research: Biogeosciences 124, 1401–1412 (2019).

50. N. G. McDowell, H. D. Adams, J. D. Bailey, M. Hess, T. E. Kolb, Homeostatic Maintenance Of Ponderosa Pine Gas Exchange In Response To Stand Density Changes. Ecological Applications 16, 1164–1182 (2006).

