## Supplemental Tables and Figures for "Experimental flooding impacts soil biogeochemistry but not aboveground vegetation in a coastal forest"

### Supplemental Information for “Experimental flooding impacts soil biogeochemistry but not aboveground vegetation in a coastal forest”

#### Table S1 - Wilcoxon results based on Friedman tests

**Table S1 - Friedman test results**

| Variable | Depth | Pre-flood | Flood | Post-flood |
|---|---|---|---|---|
| VWC (m3/m3) | 5 | ns | *** | *** |
| VWC (m3/m3) | 15 | ns | *** | *** |
| VWC (m3/m3) | 30 | ns | *** | *** |
| EC (uS/cm) | 5 | ns | *** | *** |
| EC (uS/cm) | 15 | ns | *** | *** |
| EC (uS/cm) | 30 | ns | *** | *** |
| DO (%sat) | 10 | ns | * | ** |
| DO (%sat) | 20 | ns | *** | *** |
| DO (%sat) | 30 | ns | *** | *** |
| DO (%sat) | 5 | * | ns | ** |
| Eh (mV) | 5 | ns | *** | ** |
| Eh (mV) | 15 | ns | *** | ** |
| Eh (mV) | 30 | ns | *** | *** |
| Eh (mV) | 50 | ns | * | *** |

#### Table S2 - medians and ranges of sensor measurements across flooding periods

**Table S2 - medians and ranges of sensor measurements**

| | Pre-flood | | | Flood | | | Post-flood | | |
|---|---|---|---|---|---|---|---|---|---|
| Variable | Control | Freshwater | Saltwater | Control | Freshwater | Saltwater | Control | Freshwater | Saltwater |
| VWC (m3/m3) | 0.30 (0.29-0.35) | 0.28 (0.27-0.34) | 0.27 (0.26-0.33) | 0.32 (0.30-0.35) | 0.40 (0.31-0.42) | 0.38 (0.33-0.40) | 0.30 (0.28-0.34) | 0.33 (0.28-0.42) | 0.31 (0.30-0.38) |
| EC (uS/cm) | 78.67 (50.00-121.67) | 75.67 (45.50-116.33) | 72.00 (45.00-105.67) | 76.80 (66.80-104.40) | 112.10 (77.80-153.60) | 100.20 (87.40-134.20) | 79.60 (64.20-106.00) | 1420.80 (65.80-2830.20) | 1100.30 (803.60-2156.00) |
| DO (%sat) | 92.26 (84.82-93.13) | 92.57 (84.50-94.58) | 93.39 (87.36-94.68) | 81.51 (77.24-82.69) | 22.61 (0.00-91.44) | 89.30 (0.00-91.47) | 80.79 (73.50-87.99) | 25.08 (0.23-93.45) | 77.67 (0.24-90.36) |
| Eh (mV) | 712.14 (603.91-790.04) | 717.22 (559.82-788.60) | 752.74 (570.21-790.20) | 731.06 (616.64-801.57) | 437.78 (42.99-799.04) | 438.04 (42.48-772.87) | 734.63 (665.11-765.56) | 646.23 (319.31-800.42) | 557.86 (385.59-695.66) |

#### Table S3 - Sen-Theil slopes and p-values for sap flux patterns in Figure 4

| Plot | Species | Sen-Theil slope | P-value |
|---|---|---|---|
| Freshwater | Beech | 0.000033 | ns |
| Freshwater | Red Maple | 0.000004 | ns |
| Freshwater | Tulip Poplar | -0.003920 | *** |
| Saltwater | Beech | 0.001616 | *** |
| Saltwater | Red Maple | -0.004799 | *** |
| Saltwater | Tulip Poplar | -0.008803 | *** |

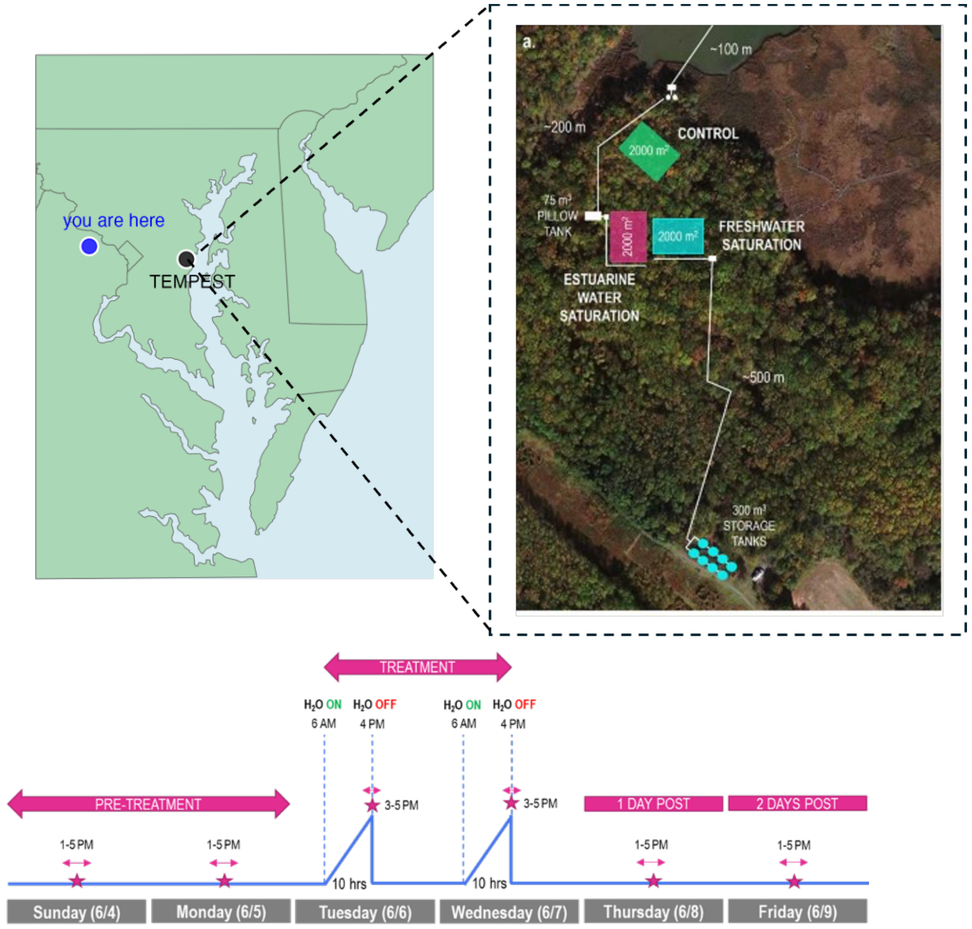


Figure S1: Map of the geographical location of the TEMPEST experiment (upper left) in Maryland, the layout of the TEMPEST plots (upper right), and the timeline of the 2023 flooding events (bottom)

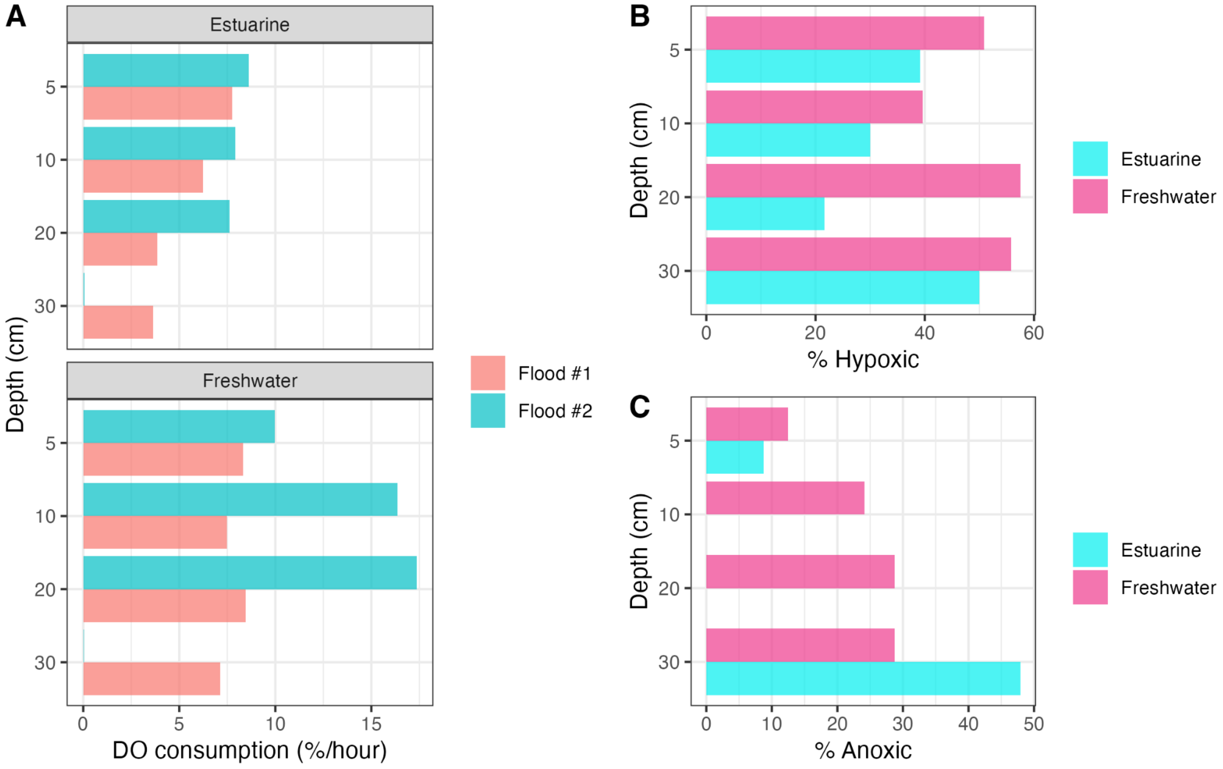


Figure S2: Statistics for dissolved oxygen (DO) measurements, including A) DO consumption rates, B) the percent of the study period shown in Figure 2A that DO levels were hypoxic (<21%) and C) the percent of the study period shown in Figure 2A that DO values were anoxic (<1%).

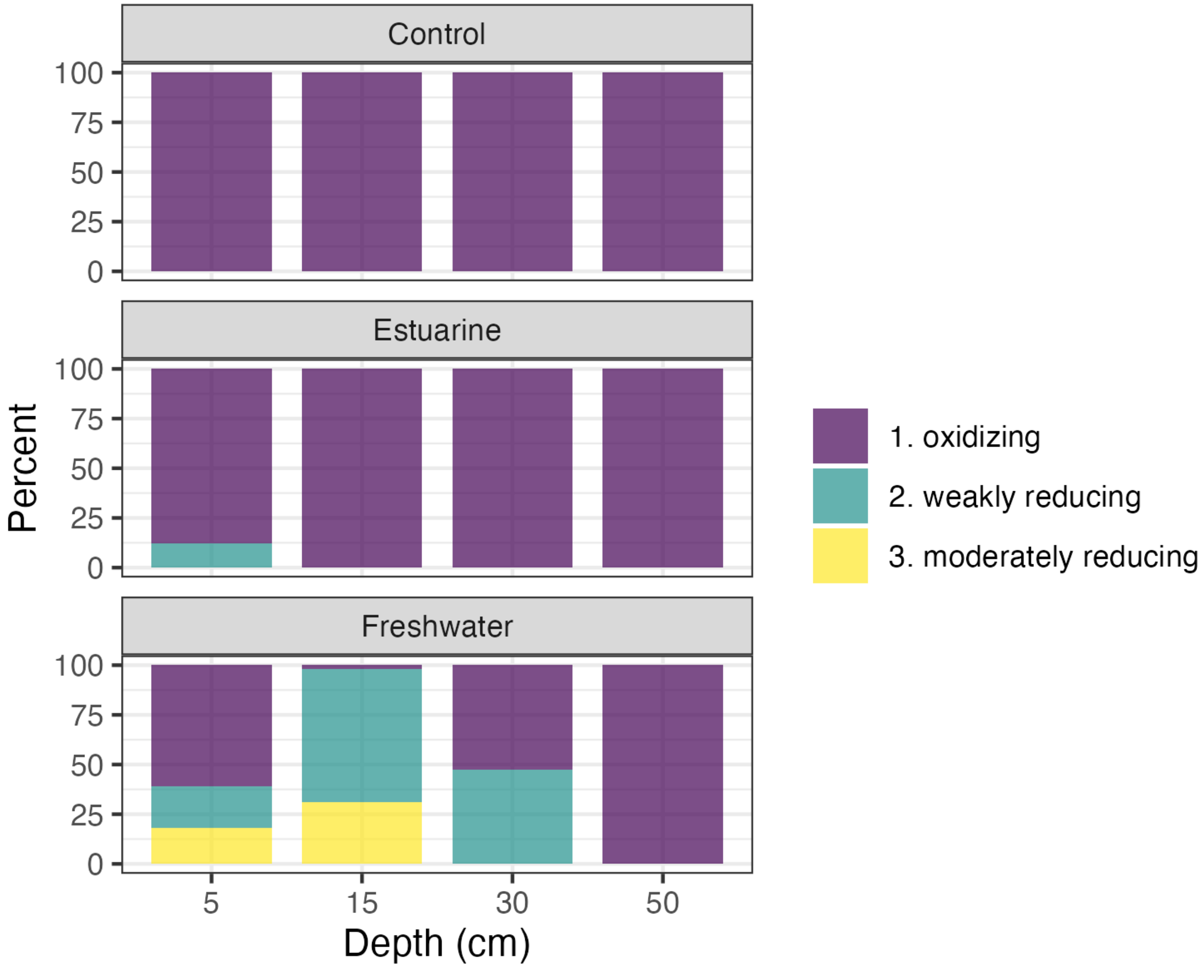


Figure S3: Redox categories defined in Machado-Silva et al. (2024) by plot for the study period show in Figure 1D.

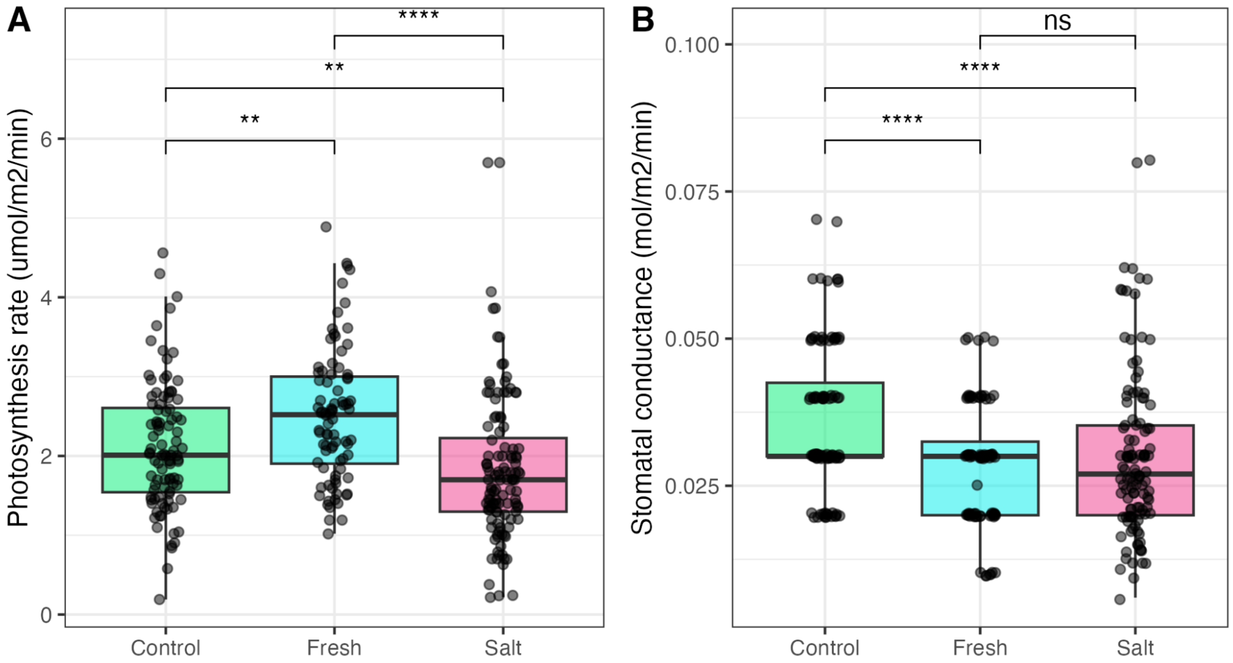


Figure S4: Measurements of intercellular leaf stress via A) photosynthesis and B) stomatal conductance, with significance based on Wilcoxon pairwise tests.

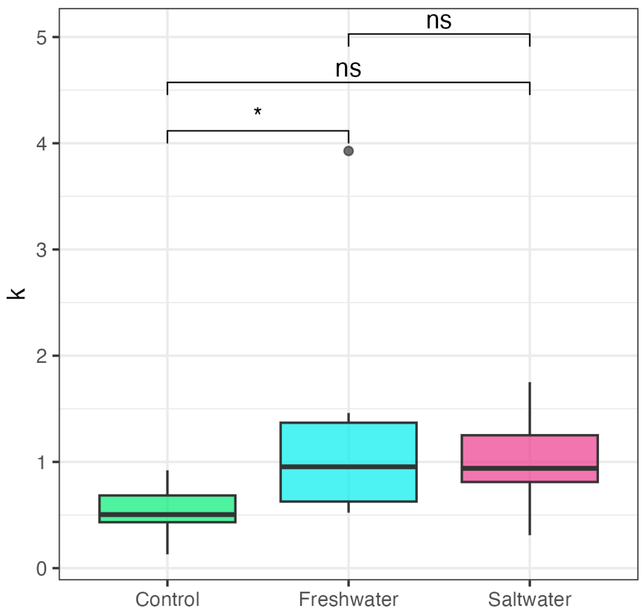


Figure S5: Belowground conductance by plot estimated as the Fick's Law constant (k) from sap flux density and water potentials, with significance based on Wilcoxon pairwise tests.
